# Stress Assessment of Vestibular Endurance Training for Civil Aviation Flight Students Based on EEG

**DOI:** 10.1101/2020.09.10.290858

**Authors:** Haixu Hu, Zhou Fang, Zhiyu Qian, Liuye Yao, Ling Tao, Bing Qin

**Affiliations:** Department of Biomedical Engineering, Nanjing University of Aeronautics and Astronautics, Nanjing, Jiangsu, 211106, China; Department of Physical Education, Nanjing University of Aeronautics and Astronautics, Nanjing, Jiangsu, 211106, China

## Abstract

**Objective:** The main goal of our study is to clarify the EEG characteristics of the stress response caused by vestibular endurance training under the real conditions.

**Methods:** Ten pilot trainees received a series of acute anti-vertigo training stimulations on the rotary ladder while recording electroencephalographic data (64 electrodes). Afterwards, the subject’s anti-vertigo ability was tested for the best performance after 1 month of training, and verifying whether it is relating to the EEG signals we collected before.

**Results:** (1) The absolute power of *α* waves in the C3 and C4 regions is same as the difference between 1 min before and 2 min after stimulation, and their activity is enhanced by stimulation. Otherwise, the activation of the C3 region after 5min of stimulation is still significant changed. (2) Discover a spearman rank correlation, the *α* waves in the C3 and C4 the greater the power change, the better the performance of the subject in the proficient stage.

**Conclusions:** C3 and C4 areas are specific brain regions of the stress response of anti-vertigo endurance training, and the absolute power of the *α* wave can be used as a parameter for identifying the degree of motion sickness (MS). The absolute power changes of *α* waves in the C3 and C4 areas are positively correlated with their anti-vertigo potential.

**Significance:** The increasing of the absolute power of *α* wave in the C3 and C4 is a manifestation of MS stress adaptability.

## Introduction

Driving an aircraft requires substantial cognitive effort and attention from operator’s brain. According to the statistics of the International Air Transport Association (IATA), more than 80% of air crashes are caused by human errors [1]. Although modern aircraft tend to be automated, pilots’ mental performance is still the most commonly contributing factor in fatal accidents worldwide. Some stimuli in real flight may lead to some psychological damage to pilots. Motion sickness (MS) is one of the main streesors for pilots to be expelled. A large number of flight accidents are caused by MS. The survey showed that the air sickness response rate is 39%, among which the pilot trainees who were expelled due to MS accounted for 55.8% of the number of those who were expelled for physical reasons [2]. Furthermore, most astronauts have also been troubled by MS problems. They often undergo MS training in the first 2-3 days in microgravity environment and the first few days after returning to earth [3]. MS is commonly used during the simulated flight training for astronauts or pilots [4]. Therefore, MS sensitivity assessment is particularly necessary for selecting pilot trainees, training quasi-pilot personnel and assessment before the restoration of the flight and after the vestibular damage is recovered [5, 6], etc. At present, traditional questionnaires, scales and physical examinations are still common methods for evaluating MS [7].

Current MS assessment methods include subjective methods and objective methods. Traditionally, researchers have used subjective questionnaires as their primary tool for evaluating pilots’ mental state [8], such as the MS Questionnaire [9, 10] and Diagnostic Criteria for Grading the Severity of Acute MS [9, 11].In addition, some researchers proposed to judge based on a specific medical history [12]. However, due to a great influence of subjective factors, the assessment accuracy is poor. With the advancement of evaluation methods and techniques, objective methods have gradually attracted attention. Commonly used objective methods including electroencephalogram (EEG) [13, 14], electrogastrogram (EGG) [15, 16], cochlear electrogram (EcochG) [17], vestibular evoked myogenic potential (VEMP) [18],heart rate variability and other physiological parameters [16].

Among objective evaluation methods, EEG recordings has been one of the most reliable contemporary methods to assess operator MS level, and they can be collected continuously without interfering with the tasks. EEG is the measurement of the electrical activity originated by the brain and recorded on the scalp surface through a net of regularly spaced electrodes. Wu (1992) found that *θ* power increased in the frontal and central areas when subjects were placed and moved in a parallel swing device [19]. Wood et al. (1994) also found increased EEG *θ* wave in the frontal areas induced by a rotating drum [20]. Yu (2010) found that the occipital, parietal, and somatosensory brain regions have significantly increased *α* and *θ* bands [21]. Chen et al (2010) believed that the *α* power of the parietal and motor areas is suppressed, and the power of the *θ* and *δ* waves of the occipital areas increased [22]. Chuang et al (2016) pointed out increasing activation of *α* and *γ* in the areas of movement, parietal and occipital areas [13]. According to the above studies, we discovered the results of MS-induced EEG power changes are inconsistent or even contradictory, such as inconsistent response areas in the cerebral cortex and conflicts of results on suppression and enhancement of brain wave signals. The reasons for this ending may be in the following three aspects, (1) the wide range of paradigms used to induce MS, (2) the EEG signal evaluation of MS level experiments are mainly based on virtual reality simulator scenes, and (3) limited to desktop device acquisition methods [7, 13, 14, 20].Therefore, there is an urgent need to supplement EEG tracking data in real and harsh environments, and analyze its characteristics to further compare and prove.

Every pilot should go through a strict training before getting their flying license. Vestibular endurance training can improve vestibular function to enhance the ability to resist MS [23]. The function of the vestibular system is to sense the three-dimensional positioning, maintain the balance of the body, and ensure that the stability of the sight is not disturbed when the head moves. Function of the vestibular system is closely related to MS and spatial disorientation in flight. 20% of flight accidents caused by spatial disorientation are caused by vestibular and visual illusions [24]. Rotary ladder training is the most commonly used vestibular endurance training and vestibular function assessment method for flying students [23]. Based on this, we collected the EEG signals of the flight students in vestibular endurance training in real time online, and combined with the final vestibular endurance performance to verify the sensitivity of the EEG assessment.

## Materials and methods

### Participants

All subjects (n = 10) volunteered to participate in the study. Besides, all of them are pilot trainees who attended the School of Flight (Nanjing University of Aeronautics and Astronautics, Nanjing, China) for aviation training. All pilots underwent a full physical examination prior to study participation. Inclusion criteria were (1) normal vision; (2) flight status at the experiment time, indicating recent good health and (3) average 7h of sleep during the night previous to the experiment without consuming alcohol before the experiment. The final sample included 9 men (one was excluded due to equipment problem), who average (±standard deviation, SD) 20.09±0.70 years age, 68.34±4.23 kg weight and 174.60±4.94 cm height. This study was approved by the Ethics Committee of Nanjing Medical University, and all subjects signed informed consent.

### Experimental design

Participants who had experienced 35±8.21 minutes rotary ladder training before firstly experienced a resting state of 5 minutes, followed by rotary ladder training stimulation, and finally rested for another 5 minutes. The EEG of all the above processes was recorded. After undergoing a rotary ladder training session for one month, the participants took the final performance test (see Study protocol and Fig 1).

**Fig 1.**
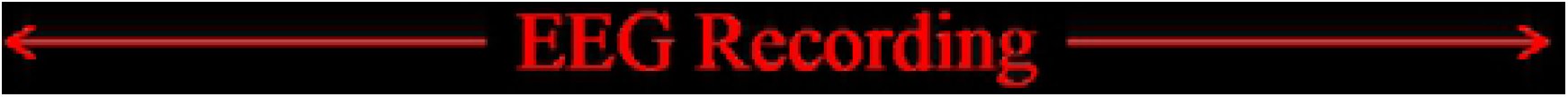
Experimental design. This fugure shows the experiment procedure we used in out study.

#### Study protocol

Rotary ladder is a basic training course for pilot trainees of Nanjing University of Aeronautics and Astronautics. Pilot trainees firstly participated in this study at the start of the rotary ladder course, i.e. in the novice stage, and all of them already have some basic rotary ladder skill. After wearing the EEG recording device on the scalp, subjects were asked to undergo a rest state for 5 minutes, while they closing their eyes, listening to the white noise through headphones and trying not to think. Then, subjects were asked to do the rotary ladder movement for 10 turns as shown in Fig 2 C. Finally, subjects immediately underwent the resting state (same as before) for another 5 minutes. since wearing EEG equipment would take time and removing the instrument halfway would affect the stimulation effect, the whole process was recorded (see EEG recording). After the first experiment, all subjects underwent a training course of rotary ladder for one month provided by the flight school. All subjects were trained at exactly the same time and were instructed by the same coach. In addition, we try to ensure that all subjects have the same practice time.The second experiment is carried out simultaneously with the final exam, each pilot student has five exam opportunities during the exam week (in fact, subjects conducted an average of 3.5 tests), the best performance score is recorded, and take it into the final exam score. The performance score has been decided by the time on how long subjects used to complete the required number of rotary ladder rotations.

**Fig 2.**
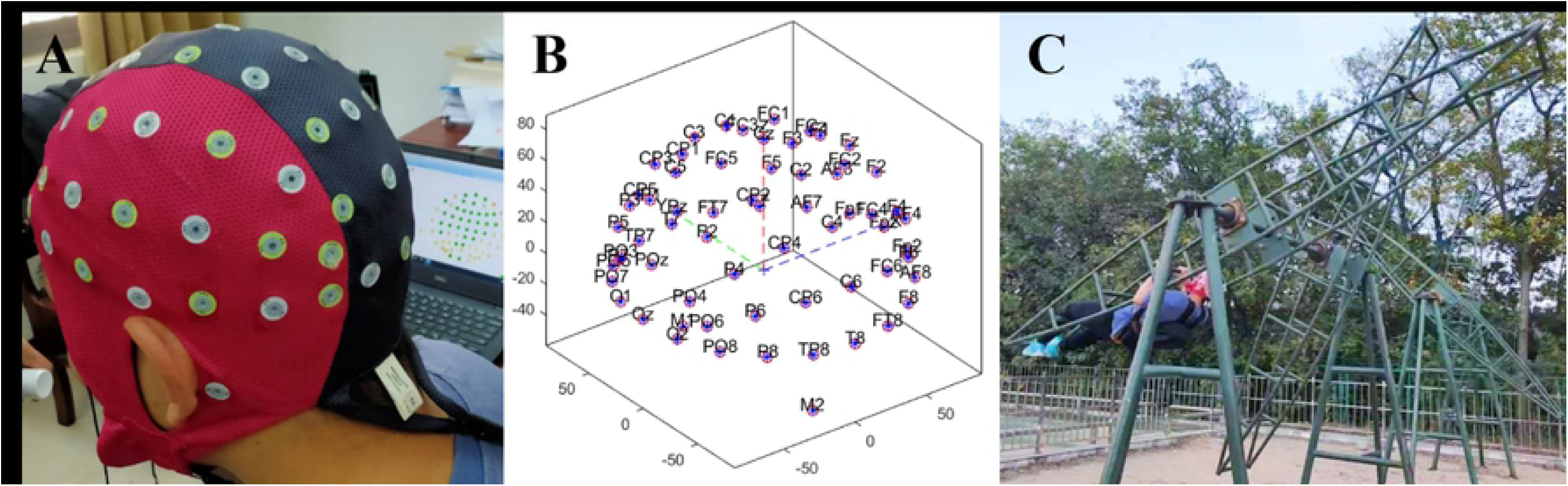
Real experiment photos and equipment structure diagram. (A) showed the diagram of device wearing. (B) showed the location of channels. (C) showed the rotary ladder movement.

#### EEG recording

We record the EEG signals using a portable EEG recorder (Waveguard Original Cap, 64 Channels, 10/10, Shielded, Tyco68). The device consisted of 3 parts (Waveguard Original Cap, computer and EEG headbox with electrodes) fastened to the back of the subject with flexible belts and was fixed stably enough to not be moved by the stimulus (i.e., artifacts from electrode movement that lead to changes in contact impedance or even the generation of a triboelectric response on the wires), as shown in Fig 2A. In order to avoid interrupting the experiment, we recorded all EEG data including the data of rotary ladder movement. The device samples data at 1000 Hz. Impedance was kept below 5 kΩ for all electrodes. We used a monopolar montage with gold cup electrodes (Natus Neurology Incorporated - Grass Products Warwick, US) at five active scalp sites: F3, F4, C3, C4, and Cz placed according to the international 10/20 system [25], and using the mastoids (A1 and A2) as references. The location of channels has been shown in Fig 2B. Ground was placed at FpZ. The channel Cz was recorded by default as an internal device requirement (internal reference). We analyzed the EEG activity of channels F3, F4, C3, and C4 (see EEG analysis). This combination was optimum for avoiding recording errors due to device vibration, electromagnetic interference, and subject movements [26].

#### EEG analysis

We analyzed EEG data using the Matlab EEGlab software package [27]. The electrode locations were based on a standard sixty-four-channel system provided by EEGlab. The acquired EEG signals were firstly inspected to remove bad EEG channels and the gross areas. To reduce the influence of artifacts, both physiological and non-physiological, in the analysis, we discarded data segments containing voltage values outside the [-100 *µ*V 100 *µ*V] interval. We intercepted the 5-minute data before and after the stimulation according to the markers and changed the data sampling rate to 256Hz. A high-pass filter with a cut-off frequency at 1 Hz was used to remove baseline-drifting artifacts. Then, a low-pass filter with a cut-off frequency of 40 Hz was applied to the signal to remove muscular artifacts and line noise.

The filtered EEG signals were decomposed into independent brain sources by independent component analysis (ICA) to separate the eye activities, such as eye blinking, lateral eye movements (LEM). The ICA algorithm can separate N source components from N channels of EEG signals and assumed the time courses of the sources are statistically independent. Therefore, The multi-channel EEG record are considered as mixtures of underlying brain sources and artificial signals. The source signals contribute to the scalp EEG signals through a fixed spatial filter. Such a spatial filter can be reflected by the rows of inverse of unmixed matrix, *W* in *u* = *Wx*, where *u* is the source matrix and *x* is the scalp-recorded EEG. The spatial filters can be plotted as the scalp topography of independent component. The ICA component decomposition result can be seen in Fig 3 A.

**Fig 3.**
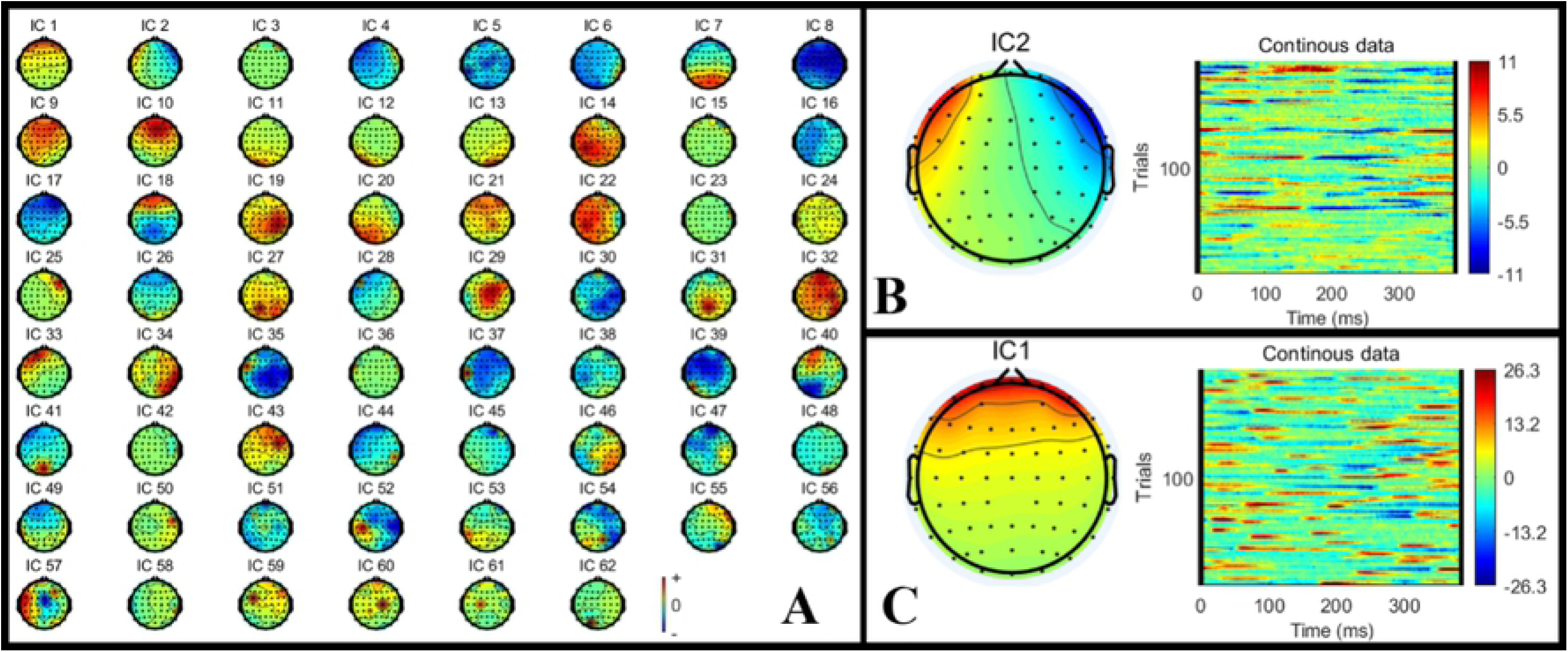
The ICA result. (A) showed the ICA component decomposition result. (B)-(C) showed the removed IC components’ topographic map and time-frequency results. (B) showed the eye blinks activities component. (C) shows the lateral eye movements component.

Eye artifacts are always present in EEG datasets. They are usually in leading positions in the component array (because they tend to be big). The eye artifact component can be identified for two reasons: (1) The smoothly decreasing EEG spectrum is typical of an eye artifact; (2) The scalp map shows a strong far-frontal projection typical of eye artifacts. Take one of the subject as an example, the removed components have been shown in Fig 3 B-C. Fig 3 B shows a typical eye blinks activities component and Fig 3 C shows a lateral eye movements component.

After removed the eye artifacts components, the EEG datasets was segmented in five epochs of 1 min each, so that the first and second epochs (b_1_, b_5_) comprised the first and the last minute of the data before the stimulation, the second, fourth and fifth epochs (a_1_, a_2_, a_5_) comprised the first, the second and the last minute of the data after the stimulation. The fast Fourier transform (FFT) implemented in the Matlab was used to perform spectral analysis and calculate power spectra for the *δ* (0.5–4 Hz), *θ* (4.0–8 Hz), *α* (8.0–13 Hz), and *β* (13–30 Hz) frequency bands. Then, we computed the average power for each frequency band, channel, and epoch.

### Statistical analysis

EEG power spectrum changes after stimulation were tested with the one-way repeated-measures analysis of variance (ANCOVA),was applied with change from baseline as the dependent variable, and treatment as independent variables. The primary focus of the analyses was the changes in outcome before and after rotary ladder training intervention. When the overall *P* value for the interaction between epochs was less than 0.05, prespecified comparisons were used to test two hypotheses: that changes in the second minute after the intervention would differ from that in the resting 1min before intervention, that changes in the fifth minute after the stimulation would differ from that in the first minute of resting stage before intervention. Spearman correlation coefficient was used to test the correlation between novice EEG power spectrum changes in specific brain regions after rotary ladder stimulation and performance of the rotary ladder test after becoming an expert, *P* value for the correlation was less than 0.05. Data for EEG power spectrum are presented as means (*±*SE) and 95% CI. Analyses were done using SPSS, version 26.0.

## Results

We monitored the EEG power spectrum of pilot’s trainee before and after the rotary ladder stimulation, in relationship to the final vestibular endurance performance.

### Effects of rotary ladder movement on EEG activity

In order to ensure that the C3 and C4 regions have been activated by the rotary ladder stimulation, we analyzed the EEG power spectrum before and after the stimulation which has been shown in Fig 4 A-D.

**Fig 4.**
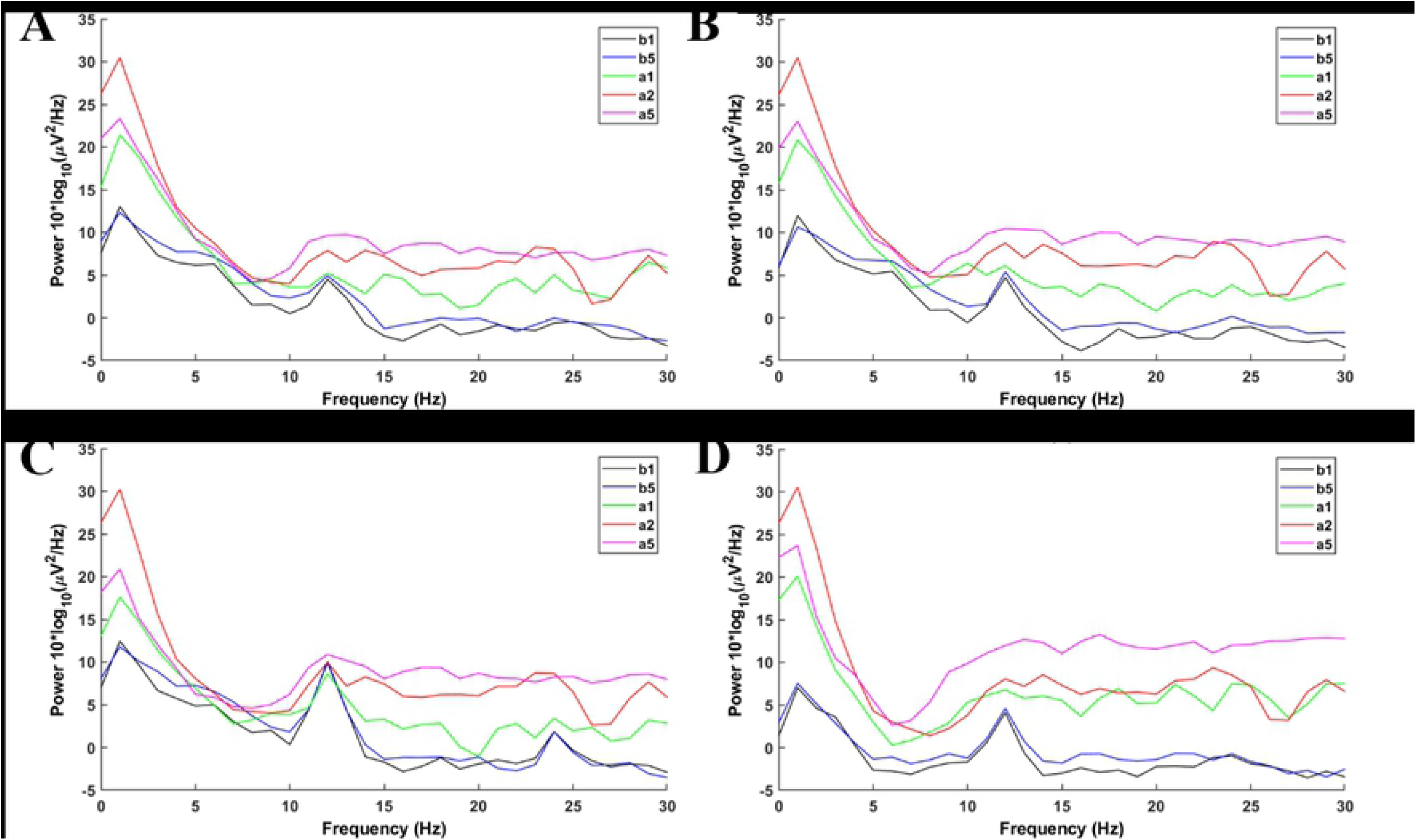
Power spectra. The power spectra of b_1_ are shown as black lines, the power spectra of b_5_ are shown as blue lines, the power spectra of a_1_ are shown as green lines, the power spectra of a_2_ are shown as red lines, the power spectra of a_5_ are shown as pink lines. (A) shows the power spectra of F3 area, (B) shows the power spectra of F4 area, (C) shows the power spectra of C3 area, (D) shows the power spectra of C4 area.

Furthermore, we analyzed the EEG power spectrum of *α* band in C3 and C4 before and after the stimulation which has been shown in Table **??**.

**Table 1.** EEG power spectrum of *α* band before and after the stimulation.

|  | C3 (n=8) |  | C4 (n=8) |  |
| --- | --- | --- | --- | --- |
|  | Mean (95% CI) | SE 1 | Mean (95% CI) 1 | SE |
| $b_1$ | 36.22 (11.25, 61.20) | 10.56 | 31.78 (11.20, 52.36) | 8.70 |
| $b_5$ | 50.89 (31.06, 70.73) | 8.39 | 52.57 (30.95, 74.19) | 9.14 |
| $a_1$ | 59.19 (26.95, 91.43) | 13.63 | 52.42 (24.66, 80.18) | 12.64 |
| $a_2$ | 62.07 (30, 94.14) | 13.56 | 58.68 (28.79, 88.27) | 12.64 |
| $a_5$ | 59.03 (35.37, 82.70) | 10.00 | 54.91 (32.15, 77.66) | 9.62 |

One-way repeated measures analysis of variance was used to determine the effect of anti-vertigo endurance training on EEG. The data of C3 is not abnormal indicated by the box plot. After Shapiro-Wilk test, the data of each group obey the normal distribution (*p>*.05). According to Mauchly’S spherical hypothesis test, the dependent variable variance covariance matrix is equal, *χ*^2^ = 0.204, *P* = 0.903. The subjects’ absolute EEG powers at the first minute before the intervention (b_1_), the second minute after the intervention (a_2_) and the fifth minute after the inervention (a_5_) were 36.22*±*10.56, 62.07*±*13.56 and 59.03*±*10.00, respectively. The difference between absolute power of EEG before and after intervention is statistically significant, *F* (2, 14) = 7.584, *p<*.01. Compared with the first minute before the intervention (b_1_), the *α* absolute power of the EEG increased (25.842) significantly in the second minute after the intervention (a_2_), *p<*.05. The absolute power of EEG in the fifth minute after the intervention (a_5_) was significantly increased by 22.808 compared with the first minute before the intervention (b_1_), *p<*.05. Fig 5 shows the changes before and after.

**Fig 5.**
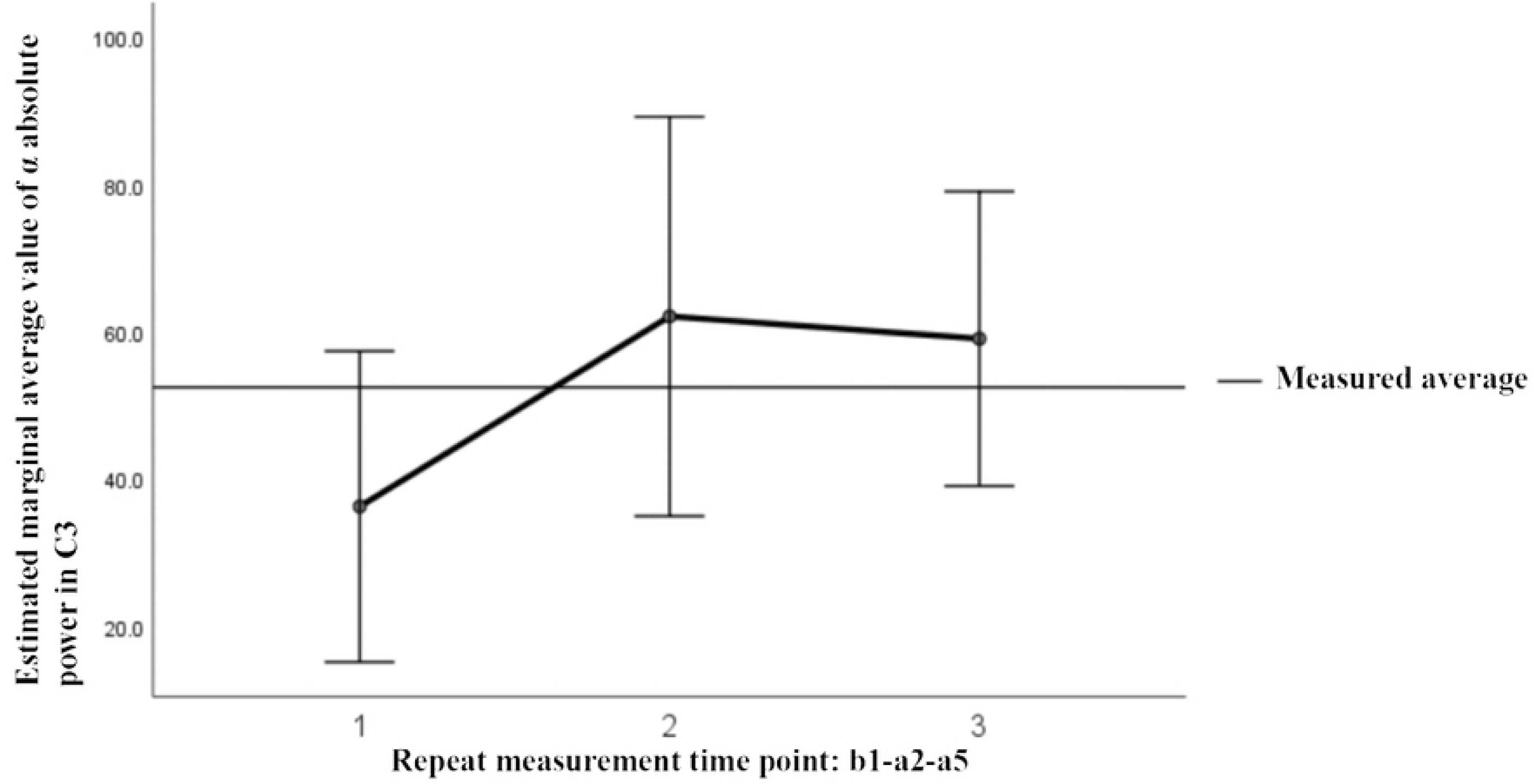
Variance analysis of repeated measurement of *α* absolute power in C3.

Similarly, C4’s Mauchly’S spherical hypothesis test indicates that the dependent variable covariance matrices are equal, *χ*^2^ = 1.931, *p* = 0.381. The subjects’ *α* absolute powers at the first minute before intervention (b_1_), the second minute and the fifth minute after intervention (a_2_ and a_5_) were 31.87*±*8.70, 58.68*±*12.64 and 54.91*±*9.62, respectively. The difference in *α* absolute power before and after intervention was statistically significant, *F* (2, 14) = 8.471, *p<*.01. The absolute power of *α* at the second minute after the intervention (a_2_) was significantly increased by 26.899, *p<*.05, compared to the first minute before the intervention (b_1_). Also, the absolute power of the *α* at the fifth minute after the intervention (a_5_) was significantly increased by 23.124 than before the intervention (b_1_), *p* = .074. Fig 6 shows the changes before and after.

**Fig 6.**
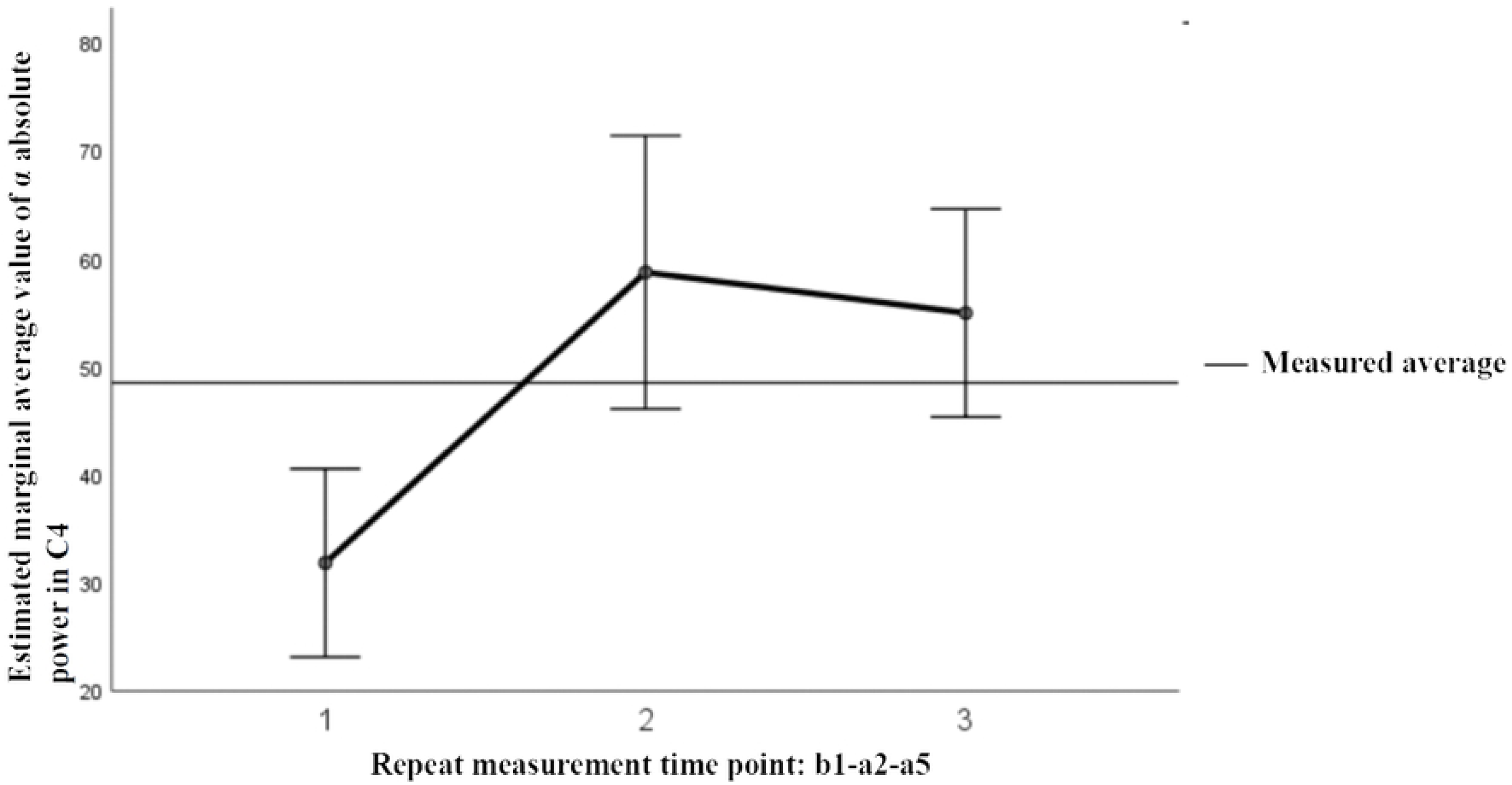
ariance analysis of repeated measurement of *α* absolute power in C4.

The results show that the absolute power of the *α* waves in the C3 and C4 regions is the same as the difference between the first minute before and the second minute after the stimulation, and their activity is enhanced by stimulation. Therefore, we choose the difference between a_2_ and b_1_ for further analysis.

### Relationship between EEG activity and vestibular endurance performance

To clarify the relationship between activation enhancement and the endurance training stimulation of the rotary ladder, we correlated the pilot trainee’s EEG response with the performance of the rotary ladder after long-term training as shown in Table 2.

**Table 2.** Correlation between expert’s final performance and novice’s EEG response of acute stimulation. Spearman correlation coefficient with the performance after long-term training, *ρ*_*X*1_ = −0:81, *p* = 0:027, *ρ*_*X*2_ = −0:81, *p* = 0:027. The performance of case 3 is 58s, which is much higher than the average value of 67.38s and rejected.

| Performance (s) | Power difference<br>of $\alpha$ in C3<br>$X_1 = T_{a2} - T_{b1}$ | Power difference<br>of $\alpha$ in C4<br>$X_2 = T_{a2} - T_{b1}$ |
| --- | --- | --- |
| 76 | 14.58 | 16.3 |
| 66 | 53.98 | 57.18 |
| 64 | 20.86 | 25.96 |
| 64 | 57.42 | 57.25 |
| 68 | 40.60 | 36.68 |
| 73 | -3.05 | 2.39 |
| 70 | 15.57 | 17.90 |

The results show that the absolute power change of the *α* waves in the C3 and C4 before and after the stimulation in the novice stage is proportional to the performance score of the proficient stage, i.e. the *α* waves in the C3 and C4 the greater the power change, the better the performance of the subject in the proficient stage, *ρ*_*X*1_ = *−*0.81, *p* = 0.027; *ρ*_*X*2_ = *−*0.81, *p* = 0.027. Therefore, the novice acute rotary ladder stimulation may be used to predict the potential skill of the rotary ladder endurance of pilot students.

## Discussion

EEG signals reflect the amounts of neurons that discharge at the same time [28]. Therefore, its time resolution is high enough to reflect the current oscillatory activity and thought to be related to the cortical resources employed for information processing [29]. Here, we examined how EEG signals reflect vestibular endurance performance. Our results indicate a differential representation of high and low score of vestibular endurance performance in the EEG signals of pilots. Pilots always need a high level of vestibular endurance that can resistant MS [5]. Previous studies have found a relationship between MS levels and EEG activity. Although these results are inconsistent, EEG is still the one of the most reliable contemporary methods to assess the MS level [22]. Increased MS level may lead to changed power in the *α* bands, i.e., the 8-12 Hz bands [22], as found here. However, little is known about the EEG response characteristics of vestibular endurance training (rotary ladder movement can cause MS) under real and harsh environments. Also, the correlation between EEG characteristics and the training plasticity of pilot trainees is even more unknown. A qualified flight selection and training needs to accurately assess the pilot trainees’ vestibular endurance performance. However, the ability of pilot candidate’s vestibular endurance is unpredictable. So, we have designed an EEG response experiment of vestibular endurance training in a real environment. We use EEG to continuously track the electrical signals of pilot students that cause MS during rotary ladder training, and objectively extract specific changes in specific brain regions. Our results indicate that the vestibular endurance training caused significant changes in *α* band wave’s power in the C3 and C4 area. Furthermore, we also found that the activations of *α* band in C3 and C4 were positively correlated with the performance of the rotary ladder after one month of systematic training. The following sections describe the details.

### Rotary ladder movement induced MS

The vestibular system refers to three pairs of perpendicular semicircular canals (anterior, posterior, and horizontal semicircular canals) and two otolithic organs (ellipsoid and balloon) in the labyrinth of the middle ear. Semicircular canals are to sense the linear acceleration, and otolithic organ is to sense the gravity stimulation. The function of the vestibular organ is to sense the three-dimensional positioning, maintain the balance of the body, and ensure that the stability of the sight is not disturbed when the head moves. Vestibular system plays an important role in motion perception, eye movement and posture control [30]. These complex functions require the integration of information about the location of the eyes, head, and external environment of the vestibule, vision, and somatosensory. This process relies on the cortex and subcortical structures, such as at the subcortical level, especially in the brain stem and cerebellum, the processing of a large amount of information from the vestibule is important to control the stability of sight and posture. In contrast, spatial orientation and motion perception also require massive processing of vestibular information at the cortical level. In the cerebral cortex of non-human primates, several multimodal sensory areas that integrate vestibular, visual, and somatic sensory signals have been described. These integrated regions can be located in specific areas of the brain [31]. Vestibular function is closely related to MS and spatial disorientation in flight [32]. Vestibular endurance training can improve the vestibular function to enhance the ability. It includes two types of training: active training and passive training. The rotary ladder training is the most commonly used vestibular endurance training for pilots and astronauts. It has been considered to relate to the vestibular endurance performance [23]. So rotary ladder trainning has been a reliable evaluation method for vestibular endurance performance [23]. Therefore, we use the time for subjects to complete the required number of rotary ladder rotations as the standard to assess their vestibular endurance capability.

### MS-related *α* band changes of parietal area

MS is associate with EEG reflection [19, 20, 22]. EEG’s *α* band (8–12 Hz) has been shown to be sensitive to sensory processing [33, 34], and has been thought to reflect the integration of cortical oscillations in multi-modal sensory systems [35]. These interactions of *α* rhythm characterize complex cortical calculations and the integration effect between multiple cortical regions. Besides, *α* EEG characteristics may be the cause of postural instability, which leads to MS [36]. The vestibular system is mainly controlled by the Parieto-Insular Vestibular Cortex (PIVC), which is located in the parietal lobe, and related to the posterior parietal cortex [31, 37–39]. PIVC is related to spatial changes in the head (vestibular), neck rotation (proprioception), and movement of visual targets [38]. The conclusion of our study that vestibular endurance training interferes with the absolute power of *α* in the parietal area C3 and C4 is consistent with the mechanism of action of the *α* band and vestibular cortex mentioned above. In comparison, some studies on the EEG characteristics of MS have quite different conclusions. For example, visually-induced MS is related to the reduction of the frontal area *θ* power and the temporal area *β* power when watching 2D and 3D movies [40]. MS is also related to increased *δ* power, decreased frontal-temporal *β* power [40]. and *α* and *γ* waves in parietal and occipital regions under virtual reality driving conditions [13]. Under simulated driving conditions, MS is associated with a decrease in *θ* power and an increase in the *δ* power of the Fz and Cz electrodes [41], and parietal area, exercise, and occipital brain regions also showed significant changes in EEG power [42]. HU et al. used optokinetic drums to induce MS and found that the *δ* power of C3 and C4 was stronger than the baseline [43]. The reason for the difference in previous research conclusions is that the characteristics of EEG cannot be faithfully reflected under virtual or simulated conditions. Yu et al. found that MS is related to the increase in energy in the *α* and *θ* bands of the parietal area, occipital area, and somatosensory brain area when riding a car under the real condition [44]. Vestibular stimulation under harsh, real conditions is greater. Reichenbach et al. found that during active vestibular exercise with higher stimulation intensity, EEG signal changes mainly occurred in the left posterior parietal cortex [45]. Our results also indicate that EEG is sensitive to the MS. Here we found a significant difference between the EEG power in the 8-12 Hz spectrum (*α* bands) before and after the rotary ladder movement in the novice stage. These results are compatible with the recently observed EEG power in relation to the MS in C3 and C4 areas. After verification, we found no significant difference during the same period of time (before or after the stimulation). These results perhaps reflected the effects of rotary ladder stimulation on EEG activities. We believe that the rotary ladder movement activates the *α* band covering C3 and C4 areas. To some extent, our research reveals the EEG characteristics of high-intensity vestibular stimulation in real environments, broadens the way humans are studied, and provides objective support for the selection and evaluation of talents such as pilots and astronauts.

### EEG reflected vestibular endurance performance

We observed a significant relationship in the vestibular endurance performance (rotary ladder performance) and EEG activities. Our results show that the absolute power change of the *α* waves in the C3 and C4 before and after the stimulation in the novice stage is proportional to the vestibular endurance performance of the proficient stage. In the other words, the *α* waves in the C3 and C4 the greater the power change, the better the performance of the subject.

Our combined results indicate that EEG could be used to evaluate a pilot’s vestibular endurance ability and potential performance in flight. These findings may also enhance our understanding of the relationship between brain activity and vertigo in complex and reality situations.

## Conclusion

The present work represents the first step towards the collection of real-time online EEG signals from flight trainees during acute regular vestibular endurance training,in addition, it can also be combined with the association of vestibular endurance performance after training to verify the sensitivity of EEG assessment. The study found that the acute anti-vertigo training triggered a stress response to the absolute power of the *α* waves in the C3 and C4 areas. The absolute power change of the *α* waves in the C3 and C4 areas before and after the intervention in the novice stage is proportional to the best movement performance (the time of the best efforts to complete the same number of rotations) of the proficient stage. That is, the greater the change in the absolute power of the *α* waves in the C3 and C4 areas, the better the performance of the pilots in proficient stage in the later training. It can be seen that the novice’s EEG response is related to the performance of the rotary ladder after long-term training. The novice’sacute rotary ladder stimulation may be used to predict the potential of the rotary ladder endurance or the ability to resist MS of the pilot students. By this way, we may complete the screening of professionally qualified personnel with special requirements for vestibular function more accurately, and early warning of operation safety, etc.

## Supporting information

**1 Data. Experimental data**. Data that recorded subjects EEG alpha power in C3 and C4 regions and their performance score.

## Author Contributions

**Haixu Hu**: Study concept and design, Statistical analysis, Experimentaldesign, Drafting the manuscript, Critical review and editing the manuscript for content. **Zhou Fang**: Data acquisition, Computational image processing, Statistical analysis, Drafting the manuscript, Critical review and editing the manuscript for content. **Zhiyu Qian**: Study concept and design, Critical review and editing the manuscript for content. **Liuye Yao**: Data acquisition, Critical review and editing the manuscript for content. LingTao:Data acquisition, Critical review and editing the manuscript for content. **Ling Tao**: Study concept and design, Critical review and editing the manuscript for content. **Bing Qin**: Data acquisition, Critical review and editing the manuscript for content.

## Acknowledgments

This work was supported by the China postdoctoral science foundation, grant number 2019M661837.

